# Visualization of subcortical structures in non-human primates in vivo by Quantitative Susceptibility Mapping at 3T MRI

**DOI:** 10.1101/2021.04.17.440277

**Authors:** Atsushi Yoshida, Frank Q Ye, David K Yu, David A Leopold, Okihide Hikosaka

## Abstract

Magnetic resonance imaging (MRI) is now an essential tool in the field of neuroscience involving non-human primates (NHP). Structural MRI scanning using T1-weighted (T1w) or T2-weighted (T2w) images provides anatomical information, particularly for experiments involving deep structures such as the basal ganglia and cerebellum. However, for certain subcortical structures, T1w and T2w images fail to reveal important anatomical details. To better visualize such structures in the macaque brain, we applied a relatively new method called quantitative susceptibility mapping (QSM), which enhances tissue contrast based on the local tissue magnetic susceptibility. To evaluate the visualization of important structures, we quantified the the contrast-to-noise ratios (CNRs) of the ventral pallidum (VP), globus pallidus external and internal segments (GPe and GPi), substantia nigra (SN), subthalamic nucleus (STN) in the basal ganglia and the dentate nucleus (DN) in the cerebellum. For these structures, the QSM method significantly increased the CNR, and thus the visibility, beyond that in either the T1w or T2w images. In addition, QSM values of some structures were correlated to the age of the macaque subjects. These results indicate that the QSM method can enable the clear identification of certain subcortical structures that are invisible in more traditional scanning sequences.

**Highlights:**

- NHP subcortical structures are challenging to see in conventional T1w and T2w images
- We applied quantitative susceptibility mapping (QSM) to identify them easily
- QSM clearly visualized basal ganglia and cerebellar nucleus of high brain iron content
- CNRs of some subcortical nucleus were significantly higher in QSM
- QSM values of several subcortical nucleus increased with age

## 1. Introduction

Magnetic resonance imaging (MRI) plays an essential role in neuroscience research involving non-human primates (NHP). For example, structural MRI scanning to obtain T1-weighted (T1w) or T2-weighted (T2w) images provides important details of brain structure that guides the penetration of microelectrodes for recording neural activity or cannulae for injecting any of a range of agents into the brain, including recent virally expressed proteins for optogenetic and chemogenetic perturbation of brain circuits (Amita et al., 2020; Bonaventura et al., 2019; El-Shamayleh & Horwitz, 2019; Eldridge et al., 2016; Maeda et al., 2020; Nagai et al., 2016;). Prior to the advent of MRI, NHP neurophysiologists relied on postmortem histological evaluation of brain sections under the microscope to determine definitively the area in which they had targeted with their recordings or injections. The capacity to check recording location using MRI has thus greatly increased the accuracy and efficiency of electrophysiology experiments in non-human primates.

Increasingly, modern viral methods utilize anatomical MRI data to identify targets for stereotaxic injection (Fredericks et al., 2020). While some targets are readily visible using standard MRI contrast, others are not. For example, it has been virtually impossible to use T1w or T2w contrasts to detect boundaries within subcortical nuclei of the basal ganglia (e.g., the globus pallidus, the substantia nigra) and the cerebellum (e.g., the dentate nucleus). This is because the signal intensities of these gray matter structures are, in fact, similar to those of the white matter around them. As a result, many functionally distinct substructures whose differential targeting may be of potential interest remain undifferentiated even in high quality MRI atlases of the macaque brain (Seidlitz et al., 2018).

Quantitative susceptibility mapping (QSM) is a relatively new MRI acquisition approach for enhancing tissue contrast (Liu et al., 2011; Wu et al., 2012; Li et al., 2011). This method exploits subtle differences in local tissue magnetic susceptibility, allowing for the visibility of otherwise hidden gray matter substructures, particularly those rich in iron. Quantitative susceptibility values across the brain are reconstructed from the MRI phase images acquired using a 3D gradient echo (GRE) sequence. Previous human studies have shown that the QSM method offers substantial visualization of internal tissue contrast for structures such as the basal ganglia, whose components vary in their level of iron deposition. For example, within the basal ganglia, this method allows for the visualization and segmentation of gray matter structures with similar T1w and T2w contrasts, such as the globus pallidus external segment (GPe), globus pallidus internal segment (GPi), subthalamic nucleus (STN), substantia nigra (SN) (Liu et al., 2013b). As a result, the QSM method has been clinically used to guide neurosurgical implantation of electrodes in the STN, a primary target in deep brain stimulation (DBS) for Parkinson’s patients (de Hollander et al., 2014; Dimov et al., 2018). While the QSM should be similarly useful to neuroscientists focusing on these and other subcortical structures in monkeys, to our knowledge this method has not been previously used in NHPs at 3T.

The present study applies the QSM acquisition protocol to the macaque brain *in vivo* to examine its feasibility and robustness in this species. The results demonstrate clear contrast images of deep gray matter structure, including the visualization of subnuclei of the basal ganglia and the cerebellum that is unclear in T1w and T2w images.

## 2. Materials and methods

### Subjects

Six adult male rhesus monkeys (Macaca mulatta), three younger monkeys (M1 (8 years and three months old, 10kg), M2 (8 years and five months old, 13kg), and M3 (8 years and five months old, 11kg)), and three older monkeys (M4 (13 years and four months old, 12kg), M5 (14 years and six months old, 14kg), and M6 (14 years and seven months old, 14kg)), participated in this study. Head post implant were present in three of the six subjects to immobilize the head. All experimental procedures followed National Institutes of Health guidelines and were approved by the Animal Care and Use Committee of the National Eye Institute.

### Animal preparation for MRI

All MR images were acquired under anesthesia to avoid image artefact by head motion. Monkey’s head was fixed in an MRI-compatible stereotaxic frame. For anesthesia, atropine (0.05 mg/kg, i.m.) was initially injected and ketamine (10 mg/kg, i.m.) and dexmedetomidine (0.01 mg/kg, i.m.) were used for induction. Additional ketamine (5 mg/kg, i.m.) and dexmedetomidine (0.01 mg/kg, i.m.) were injected for maintenance.

### Imaging Protocols and QSM reconstruction

MR imaging of all six monkeys was performed in a clinical 3T MR imaging system (MAGNETOM Prisma; Siemens Healthcare, Erlangen, Germany) using a human 15-channel human knee coil. T1w images were acquired using MPRAGE whose parameters were as follows: 0.5 mm isotropic, FOV 128 x 128 x 112 mm, matrix 256 x 256 slices per slab 224, sagittal orientation, number of averages 4, TR 2200 ms, TE 2.23 ms, TI 900 ms, flip angle 8. T2w images were acquired using SPACE (Mugler et al., 2000) with the following parameters: 0.5 mm isotropic, FOV 128 x 128 x 112 mm, matrix 256 x 256, slice per slab 224, TR 3200 ms, TE 562 ms, number of averages 2. QSM was obtained with a 3D multi-echo gradient-echo (GRE) sequence with TR 50 ms, TE: 3.65/10.11/16.73/23.35/29.97/36.59/43.21 ms, bandwidth 280 Hz/pixel, flip angle 15, FOV 128 x 128 x 57.6 mm, matrix 320 x 320 x 144, number of averages 1.

QSM images were reconstructed from the phase images from 3D GRE with multiple echo, as described previously in detail (Liu et al., 2015; Wang et al., 2015) and summarized in Fig. 1A. This reconstruction process consisted of three main steps: (1) *Unwrapping of the phase images*. This step was necessary because any angle of proton phase shift lying outside the range between −π and π is folded back in the image, and this folding was abundant in raw phase images with longer TE. (2) *High-pass filtering of phase image*. This step compensated for the dominating susceptibility influence of the tissue-air interface (‘background phase’) and focused instead on local tissue phase images. (3) *Dipole deconvolution*. This step was performed to map the magnetic susceptibility in each voxel. Many algorithms to solve this dipole inversion problem have been developed (Li et al., 2011; Liu et al., 2009; Liu et al., 2012; Tang et al., 2013; Wharton et al., 2010). In this study, we used the Morphology Enabled Dipole Inversion (MEDI) method (Bollmann et al., 2019; Liu et al., 2011; Liu et al., 2012; Liu et al., 2013a; Liu et al., 2018). These three steps were performed by using the MEDI toolbox on MATLAB2019 (Liu et al., 2012, http://pre.weill.cornell.edu/mri/pages/qsm.html).

**Figure 1.**
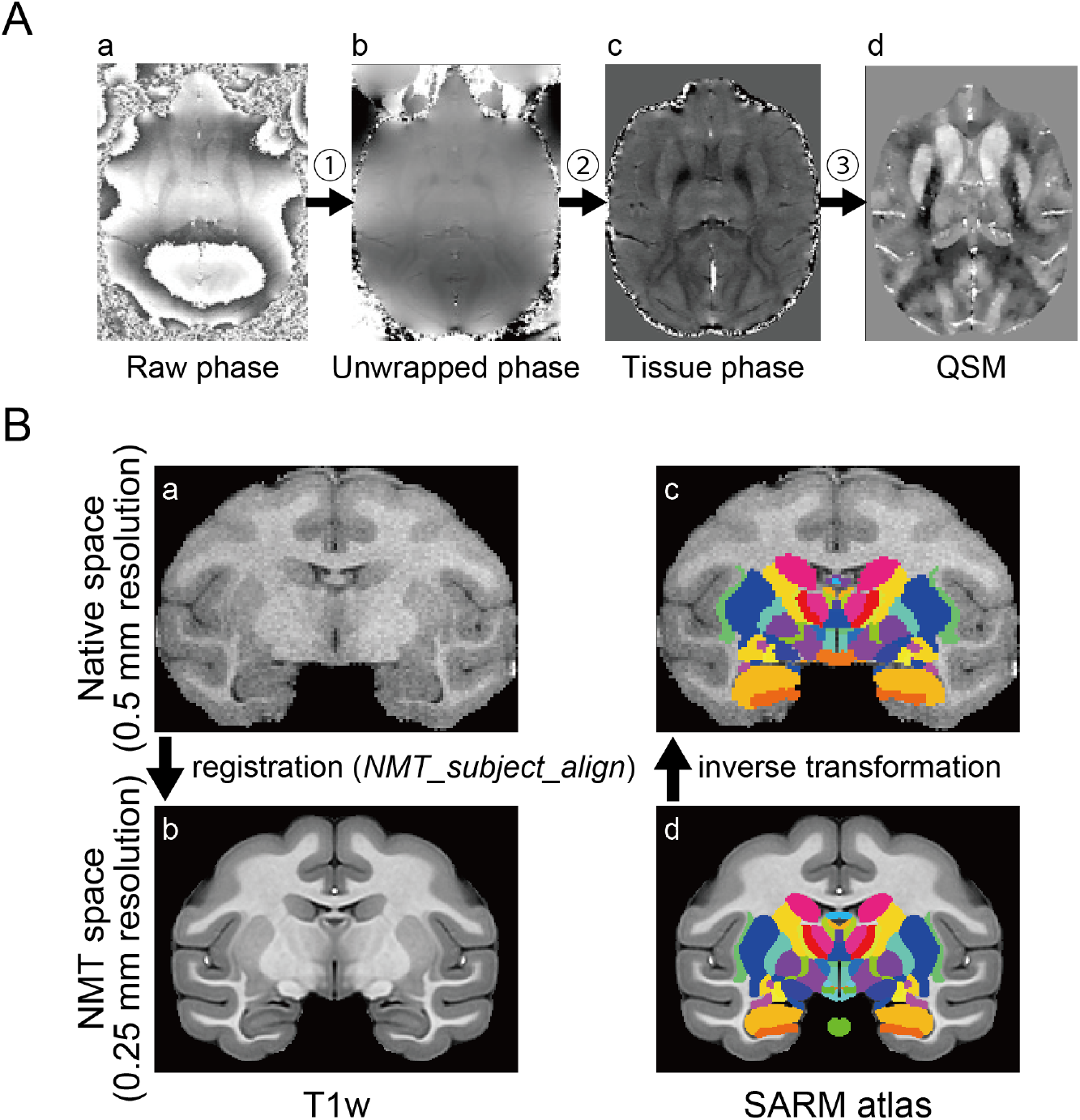
Imaging preprocesses. (A) Overview of QSM processing. QSM is reconstructed from 3D gradient echo phase images with different TE. There are three main steps in the processing from the raw phase image to QSM; ➀ unwrapping ➁ background phase removal by high-pass filtering, ➂ inverse problem solution by performing a dipole deconvolution. (B) Automatic ROI creation using SARM atlas. SARM atlas in NMT space was inversely transformed into each subject’s space (from (d) to (c)) using output from alignment of a subject’s T1w image to NMT (from (a) to (b)).

### Quantitative image evaluation

To evaluate the visibility of the ventral pallidum (VP), globus pallidus external segment (GPe), globus pallidus internal segment (GPi), substantia nigra (SN), subthalamic nucleus (STN) in the basal ganglia, and the dentate nucleus (DN) in the cerebellum in T1w, T2w, and QSM, the contrast-to-noise ratios (CNRs) were calculated by the following formula: CNR = |I_ROI_ – I_wm_|/σ_wm_, where I_ROI_ is the average signal intensity of the region of interest (ROI) and I_wm_ is the average signal intensity of white matter near ROI (Dimov et al., 2018). σ_wm_ represents the noise measurement calculated as the standard deviation of I_wm_. First of all, to create ROIs semi-automatically, each subject’s T1w image was aligned to the standard NIMH macaque template (NMT v2.0) using analysis pipeline (*NMT_subject_align*) with software AFNI (Cox 1996; Jung et al., 2021; Seidlitz et al., 2018) (Fig. 1B (a), (b)). Secondly, the subcortical atlas of the rhesus macaque (SARM) for NMT (Hartig et al., 2021) was inversely transformed to the subject T1w image (Fig. 1B (c), (d)) by AFNI’s *3dNwarpApply* with output files from NMT_subject_align pipeline (inverse of linear transformation matrix and non-linear warp field). Finally, if the modification of ROIs was needed, we edited it manually, referring to the QSM, T1w, and T2w images. Afterward, these masks were applied to the T2w and QSM images for calculating the CNRs after resampled the voxel size of the QSM from 0.4 mm to 0.5 mm as well as the T1w and T2w. We used the left hemisphere for determining ROIs. For measuring the CNR of ROIs, we defined as I_wm_ the ipsilateral anterior commissure (AC) (for the VP in Fig. 2), the internal capsule (IC) (for the GPe and GPi in Fig. 3), the optic tract (for the SN and STN in Fig. 4), the part of the ipsilateral middle cerebellar peduncle (for the DN in Fig. 5).

**Figure 2.**
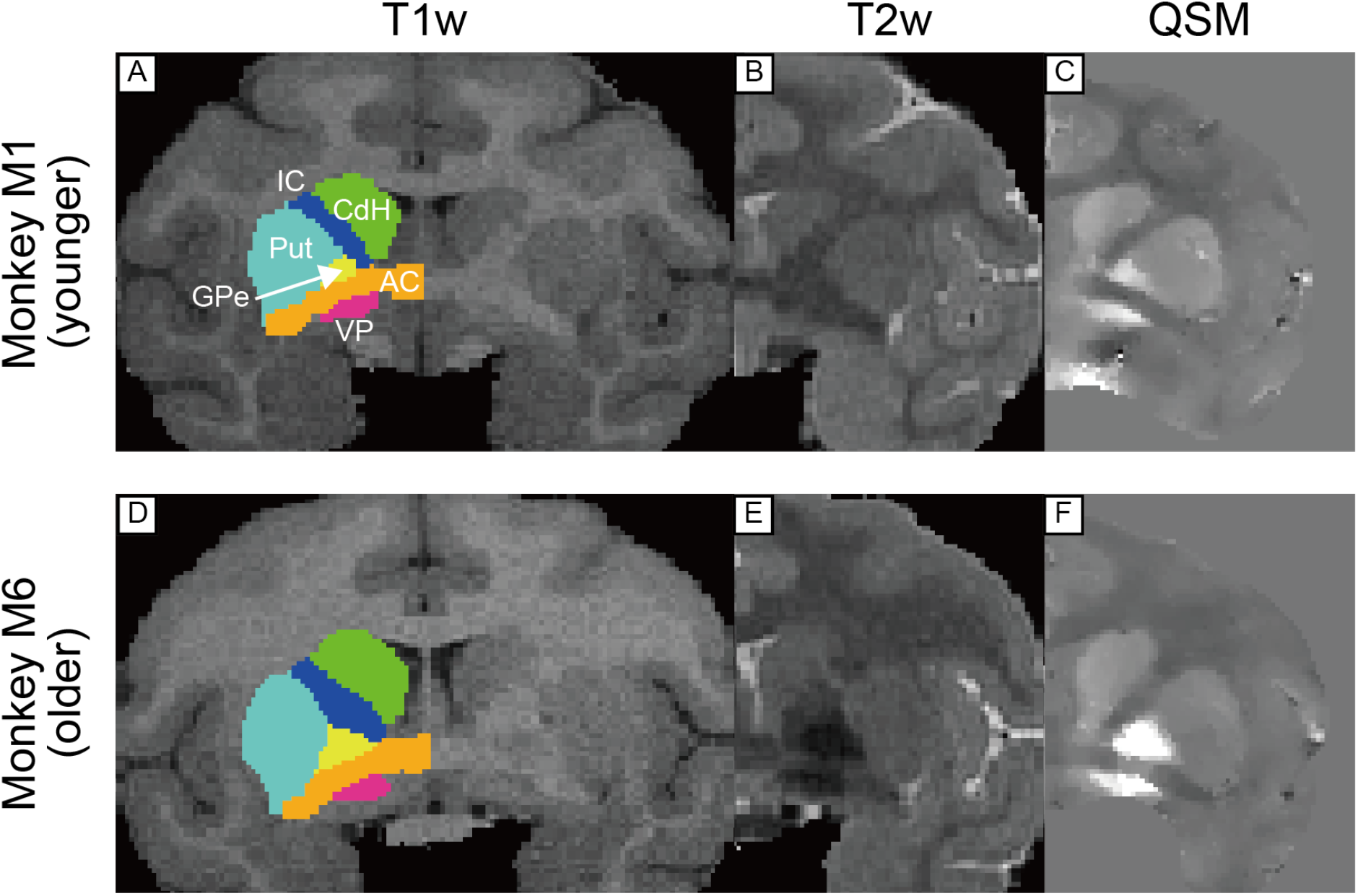
T1w, T2w, and QSM coronal images including anterior commissure (AC) in a younger ((A), (B), and (C)) and older monkeys ((D), (E), and (F)). Colored drawings in left hemisphere in T1w images indicate ROIs created from SARM atlas. These were used for calculations of CNR and mean QSM values. Abbreviations: AC, anterior commissure. CdH, head of caudate nucleus. GPe, globus pallidus external segment. IC, internal capsule. Put, putamen. VP, ventral pallidum.

**Figure 3.**
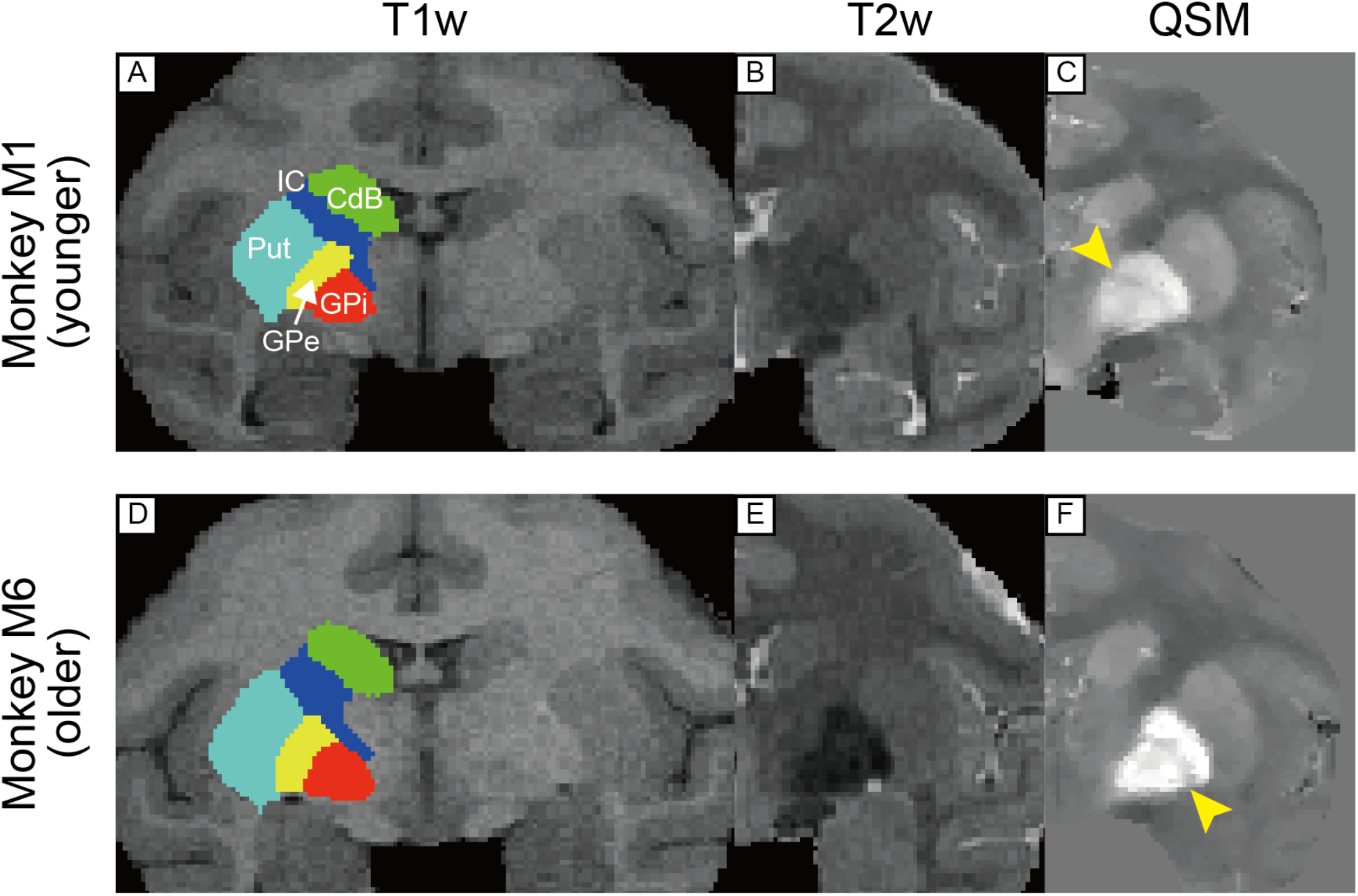
T1w, T2w, and QSM coronal images including borders of GPe and GPi in a younger ((A), (B), and (C)) and older monkeys ((D), (E), and (F)). Yellow arrow head in (C) and (F) indicates the medial medullary lamina. Abbreviations: CdB, body of caudate nucleus. GPe, globus pallidus external segment. GPi, globus pallidus internal segment. IC, internal capsule. Put, putamen.

**Figure 4.**
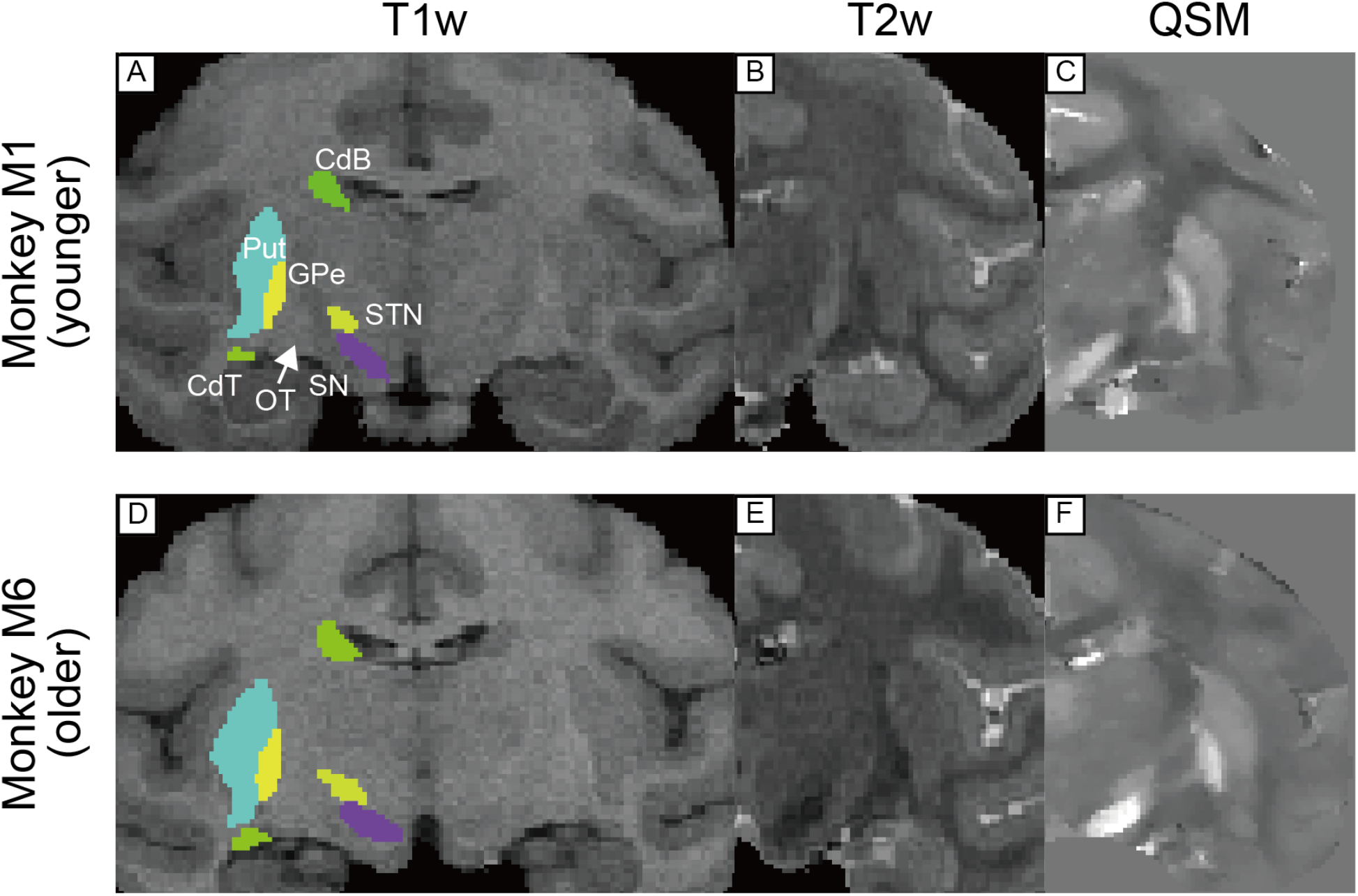
T1w, T2w, and QSM coronal images including substantia nigra in a younger ((A), (B), and (C)) and older monkeys ((D), (E), and (F)). Abbreviations: CdB, body of caudate nucleus. CdT, tail of caudate nucleus. GPe, globus pallidus external segment. OT, optic tract. Put, putamen. SN, substantia nigra. STN, subthalamic nucleus.

**Figure 5.**
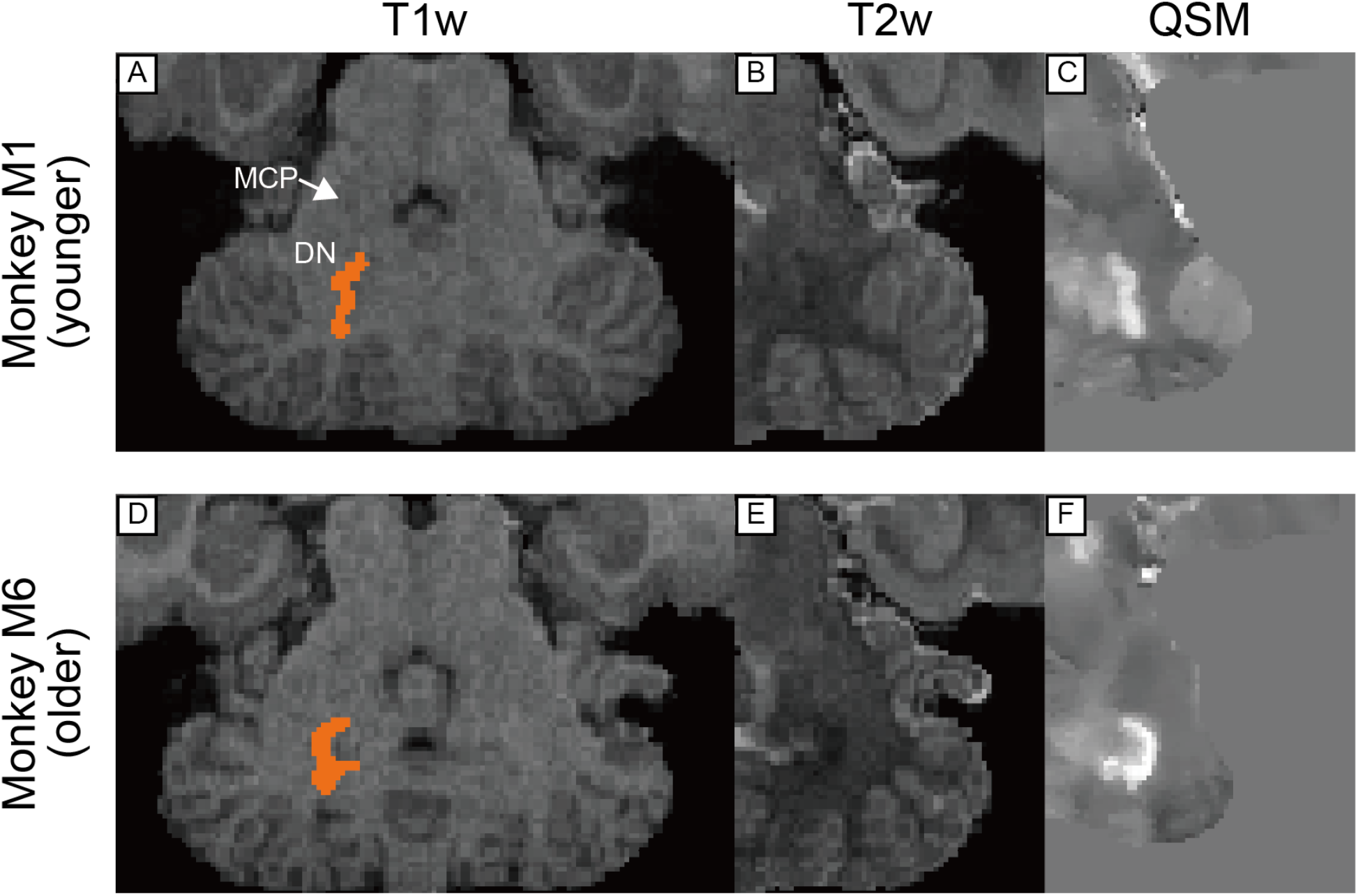
T1w, T2w, and QSM transverse images including dentate nucleus in cerebellum in a younger ((A), (B), and (C)) and older monkeys ((D), (E), and (F)). Abbreviations: DN, dentate nucleus. MCP, middle cerebellar peduncle.

### Statistical analysis

Kruskal-Wallis tests were performed to compare the differences in the CNRs among T1w, T2w, and QSM and in the subcortical structures (VP, GPe, GPi, SN, STN, and DN) followed by the post hoc multiple comparisons (Dunn-Sidak test) were executed. To investigate whether the QSM values in each subcortical structure were correlated with monkeys’ age, we also assessed Spearman’s rank correlation test between monkeys’ age and the mean QSM values in each ROIs. For multiple comparisons, we applied the Benjamini-Hochberg procedure controlling at a significance level of α = 0.05 (Benjamini et al, 1995). These statistical analyses were performed using MATLAB Statistics and Machine Learning Toolbox.

## 3. Results

### Visualization of subcortical structures

Figs. 2, 3, 4, and 5 show comparisons of subcortical structures among T1w, T2w, and QSM for several subcortical regions of the brain marked by low contrast in conventional anatomical MRI scans. The upper panel in each figure shows MR images from one of the younger monkeys (monkey M1, 8 y/o), while the lower panel illustrates images from one of the older monkeys (monkey M6, 14 y/o). In each case, QSM highlighted certain substructures as having higher susceptibility, reflecting a higher level of iron content than neighboring areas. For example, QSM applied to the pallidal structure (Fig. 2C) shows distinct boundaries among the VP, GPe, and AC, in contrast to the conventional images where the boundaries were indistinct because the structures have similar T1w and T2w values. Fig. 3 illustrates the value of QSM in identifying the boundary of another portion of the globus pallidus, namely the medial medullary lamina between the GPe and GPi (indicated by yellow-colored arrowhead in Fig.3C and F), also subtle or absent in T1w and T2w scans. In Figure 4, the QSM further highlights the substantial nigra (SN) of the midbrain, and the nearby GPe components of the globus pallidus. Although the SN was detectable in QSM from both younger and older monkeys, the subthalamic nucleus (STN) was subtle even in QSM from the older monkey (Fig.4F). QSM also illustrated the position and detailed shape of the dentate nucleus in the cerebellum (Fig. 5), which was nearly invisible in T1w and T2w images. This marked difference probably stems from the elevated iron deposition in the dentate nucleus compared to other cerebellar nuclei, such as the nucleus interpositus and nucleus medialis, whose hyperintensity is observed to be less.

### Quantitative Results

To quantitatively evaluate visualization of subcortical structures in QSM, T1w, and T2w, we measured the CNRs of them in each MR image and performed statistical tests. Figure 6 presents the results of comparing the CNRs. The Kruskal-Wallis test and post hoc test revealed that the CNRs of all 6 subcortical structures in QSM were significantly higher than those in T1w and/or T2w (*P* < 0.05, Dunn-Sidak). We also assessed the regression analysis (Pearson’s rank correlation test) to investigate whether the mean QSM value of each ROI correlated with subjects’ ages (Fig. 7). The QSM values of the VP, GPe, GPi were significantly correlated with ages corrected by Benjamini-Hochberg procedure.

**Figure 6.**
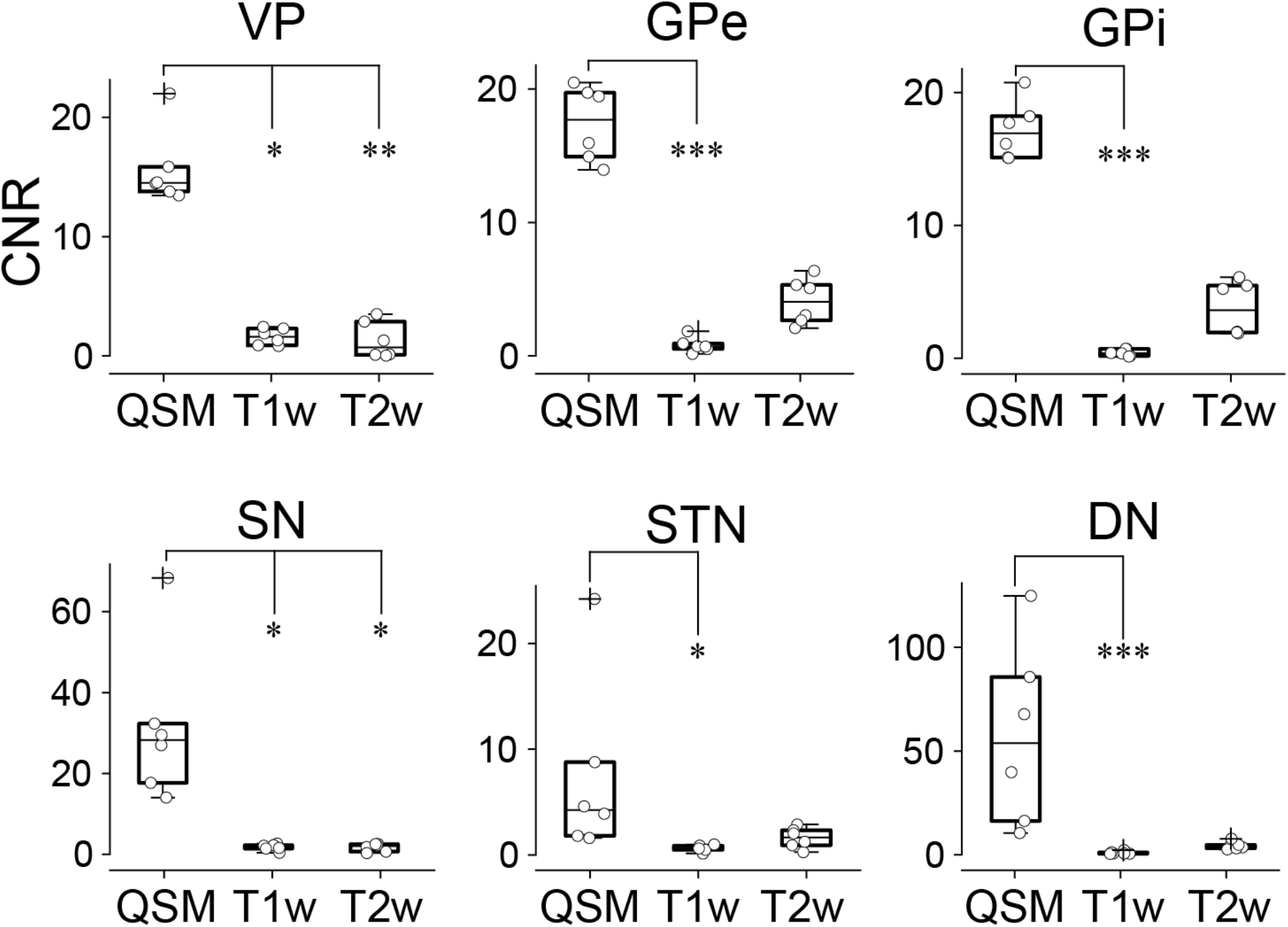
Comparison of CNR among subcortical regions in T1w, T2w, and QSM. In the box plots, upper and lower lines indicate the 75^th^ and 25^th^ percentile respectively, and dot line shows the median. Whiskers above and below the box represent the 10^th^ and 90^th^ percentiles. The cross sign indicates the outliers and the circle the CNR value of each data. Differences of CNR among images were assessed using Kruskal-Wallis test and post hoc test (Dunn-Sidak) (\**P* < 0.05, ** *P* < 0.01, \*\*\**P* < 0.001). Abbreviations: DN, dentate nucleus. GPe, globus pallidus external segment. GPi, globus pallidus internal segment. SN, substantia nigra. STN, subthalamic nucleus. VP, ventral pallidum.

**Figure 7.**
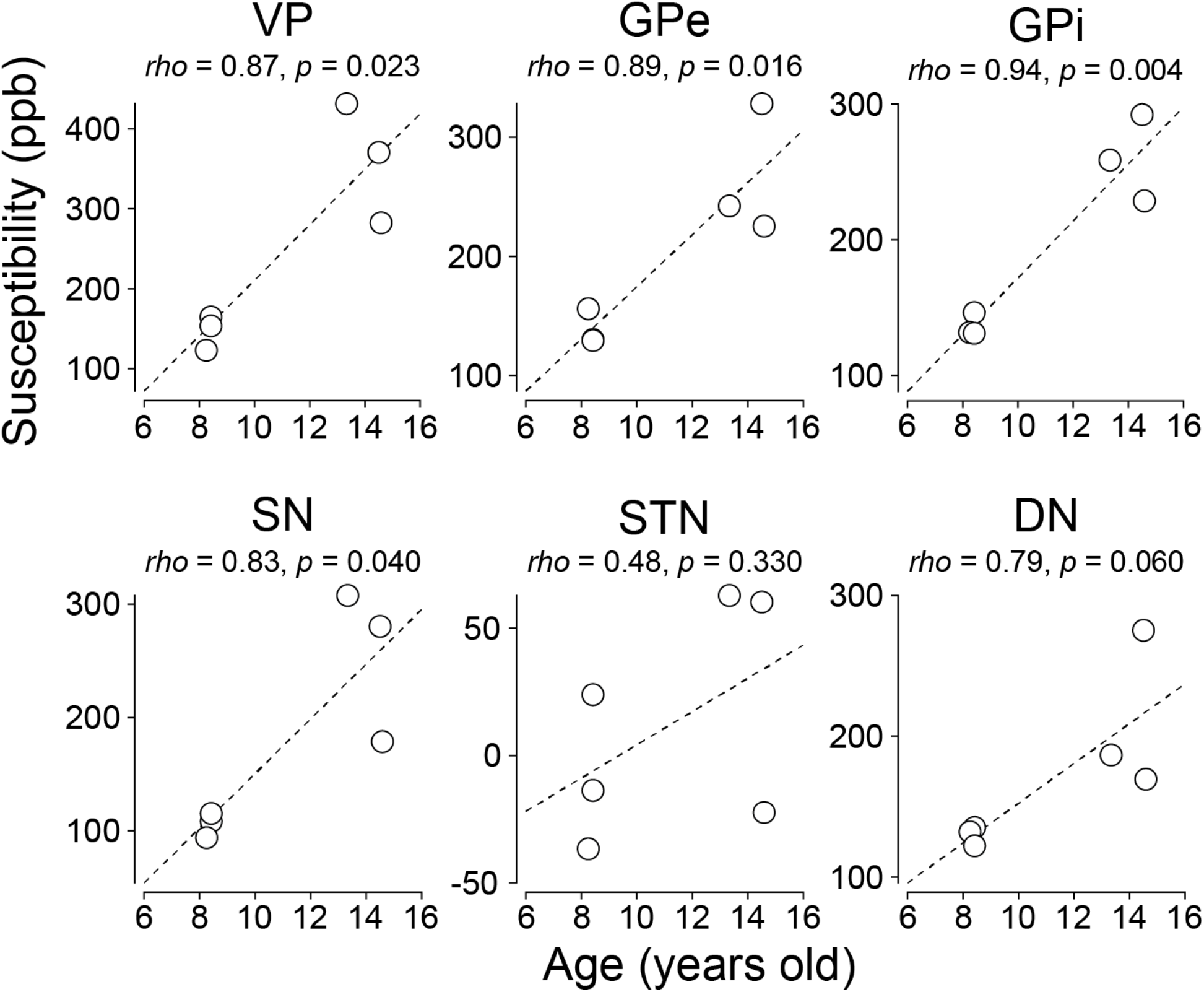
Correlation between monkeys’ ages and mean QSM values of subcortical areas. Each circle indicates the mean QSM value of individual subcortical ROIs and the dash line represents regression slope.

## 4. Discussion

Our results indicate that quantitative susceptibility mapping in the macaque offers a marked improvement in the visualization of certain subcortical structures, significantly heightening the CNRs over T1w and/or T2w images. Most notably, QSM enabled a clear and efficient differentiation of the pallidum (VP, GPe, and GPi), substantia nigra, and cerebellar nucleus (DN) from the white matter around them. Thus, this anatomical scanning approach may be advantageous for neuroscientists targeting these and nearby structures for the recording of neuronal activity or the injection of viral vectors or other agents. To our knowledge, this is the first study for the feasibility of QSM in the macaque brain in vivo at 3T MRI, with only one previous article applying QSM to the macaque brain at 9.4T (Wen et al., 2019).

QSM is based on the local measurement of magnetic susceptibility tissue, which strongly reflects the concentration of paramagnetic unpaired electrons. This sensitivity makes QSM able to identify and delineate the boundaries of the structures prone to iron deposition as ferritin, such as the nuclei in the basal ganglia (the VP, GPe, GPi, and SN) and the cerebellum (DN) (Schweser et al., 2011; Schafer et al., 2012). Iron deposition in these structures occurs from a young age in humans (Bilgic et al., 2012; Keuken et al., 2017; Li et al., 2014; Persson et al., 2015) and probably also in macaque monkeys. Actually, the CNRs of QSM, even in younger monkeys around 8 years old, were higher, and the QSM values in subcortical structures were positively correlated with ages.

The improved CNRs of the subcortical structures compared to T1w and T2w, QSM allows macaque researchers to investigate the substructure of deep subcortical regions in a way that has not previously been possible using anatomical MRI methods. This method may be particularly useful for neuroscientists in NHP who investigate the functions of the basal ganglia and the cerebellum. Clear visualization of the structures in QSM can increase the efficiency of researchers by decreasing the number of electrode or needle penetrations required for neuronal recording the injection of chemical substances such as pharmacological agents or viral vectors for genetical manipulations. Several human neurosurgery researchers have recently reported that QSM is useful for surgeries to put electrodes for deep brain stimulation (DBS) for Parkinson’s disease or dystonia patients. Similar to the work shown here, these studies showed that QSM at 3T MRI (Dimov et al., 2018; Li et al., 2020; Wei et al., 2019) and 7T MRI (de Hollander et al., 2014) visualized targets of the DBS (the STN, GPi, and centromedian thalamic nucleus). For macaque research, this method, combined with T1w images for the grid in the neuronal recording chamber, will allow us to achieve more precise targeting in focusing on the subcortical regions. The adoption of QSM may also help avoid the need to euthanize monkeys merely to check the position of electrodes or injections in histological sections.

One possible application of this method is the incorporation of QSM contrast into new macaque MRI brain atlases, such as the National Institute of Mental Health Macaque Template (NMT), which is a high-quality contrast made from T1w images of 31 macaque monkeys (Jung B et al., 2021; Seidlitz et al., 2018). Whereas the NMT shows clear structures in the basal ganglia, the T1w images, particularly in younger monkeys, do not provide sufficient contrast to segment the substantia nigra (SN). This is why one can’t get precise alignment between the standard template and the native structure image. In fact, the results of aligning the SN region in the SARM atlas to the native subject’s T1w images were not accurate (all SN ROIs were out of alignment in the superior direction (Fig. S1)), and thus we needed to manually modify ROIs for the SN of all subjects. To use QSM image for registration to NMT and SARM will provide precisely the SN discrimination, and it help us to enable the more accurate analysis, in which, for example, it is allowed for analysis of functional MRI (fMRI) data (task fMRI or resting-state fMRI) to use more accurate seed or ROI. QSM in this study also clearly illustrated the DN in the cerebellum. The DN receives projections mainly from Crus I and Crus II in the cerebellar hemisphere, which are assumed to be involved in the cognitive functions (Bostan et al, 2018; Ito 2008; Middleton et al., 2001; Strick et al., 2009). Many researchers in NHP have been interested in investigating the cognitive functions of the DN (Ashmore et al., 2013; Kunimatsu et al., 2018; Lu et al., 1998; Ohmae et al., 2013). Thus, QSM may be useful not only for neuronal recording from the DN but also for more precision seed analysis of the DN in the fMRI research in NHP.

Finally, there are a few limitations of this method that should be stated explicitly. First, QSM is highly sensitive to tissue with substances that induce susceptibility change. After penetration of electrode or needle, hemosiderin deposit will happen naturally in the affected brain tissue. Because hemosiderin includes iron, QSM illustrates hemosiderin deposit as hyperintensity. Thus, it may not be easy to see the accurate borders of the subcortical structures in QSM after several electrode or needle penetrations. Therefore, it may be better to carry out QSM scanning before any penetrations or early in the experiment. Second, iron deposition in the subcortical structures starts during adolescence in humans. Thus, the QSM method does not show hyperintensity in these structures in childhood and may therefore not be useful to guide experiments in infant monkeys (Bilgic et al., 2012; Li et al., 2014; Persson et al., 2015).

In summary, our results suggest that QSM visualized the subcortical structures more clearly than the conventional T1w and T2w. It may be useful for a range of neuroscience applications in NHP that benefit from the clear visualization of structural divisions among subcortical regions.

## Data Availability

MR images data related to this publication will be available upon reasonable request from the corresponding author (A.Y.).

## Supplementary Figure

**Supplementary Figure 1.**
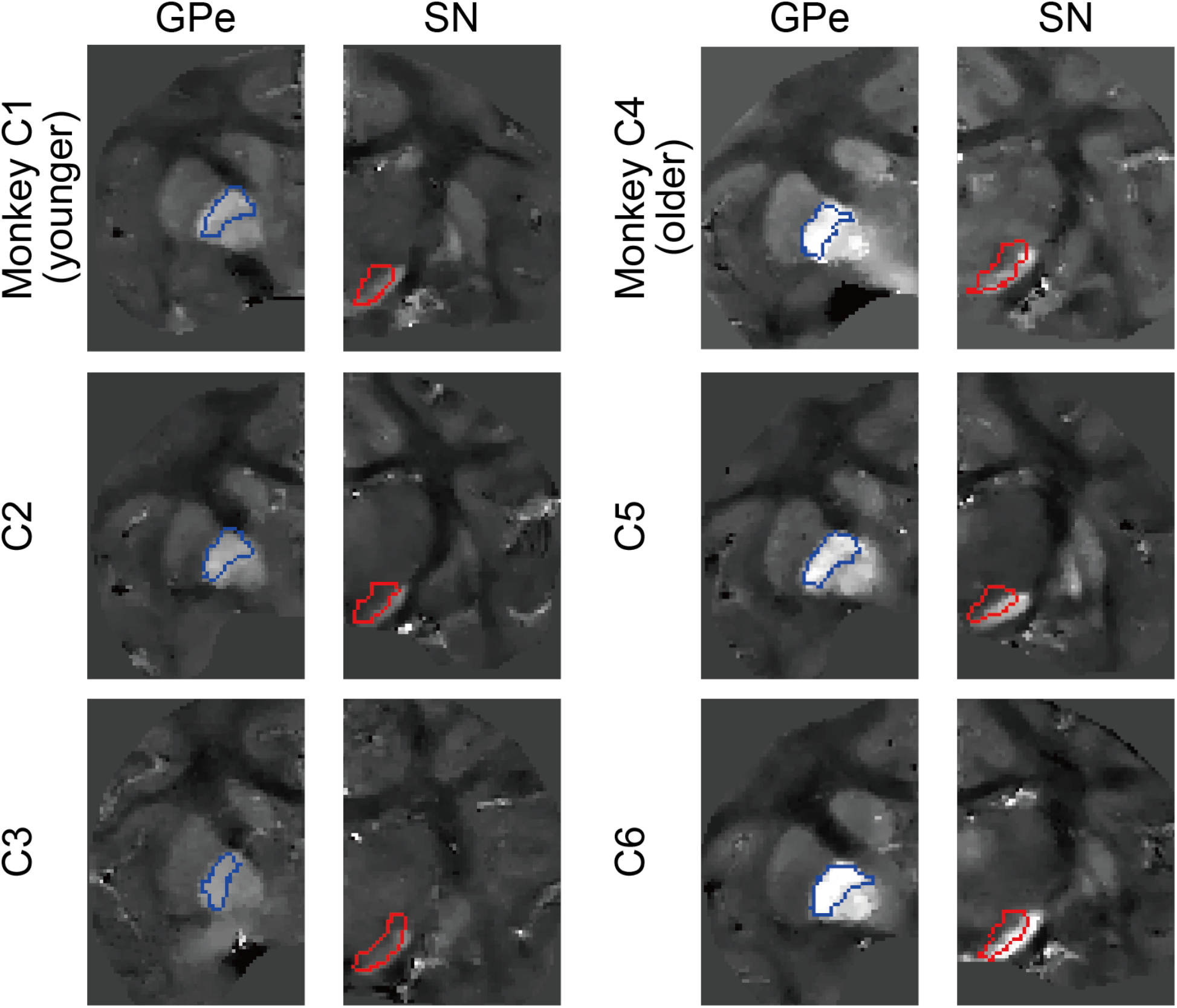
Comparison of registered GPe and SN ROIs from the SARM atlas with QSM images of all six monkeys (left hemisphere including the GPe and right hemisphere with the SN). Blue- and red-colored enclosed areas imply the edges of the GPe and SN ROIs created from the SARM atlas registration, respectively.

## Acknowledgments

This research was supported by the Intramural Research Program at the National Institutes of Health, National Eye Institute, and JSPS KAKENHI Grant (Grant-in-Aid for Fostering Joint International Research (A), 17KK0180). MRI scanning was carried out in the Neurophysiology Imaging Facility Core (National Institute of Mental Health, National Institute of Neurological Disorders and Stroke, and National Eye Institute). We thank D. Parker, A. Lopez, D. O’Brien, V.L. McLean, I. Bunea, G. Tansey, M.K. Smith, A.M. Nichols, D. Yochelson, J.W. McClurkin, A.V. Hays, and J. Fuller-Deets for technical assistance.

## Abbreviation

AC: anterior commissure
CNR: contrast-to-noise ratio
CdB: body of caudate nucleus
CdH: head of caudate nucleus
CdT: tail of caudate nucleus
DBS: deep brain stimulation
DN: dentate nucleus
GPe: globus pallidus external segment
GPi: globus pallidus internal segment
IC: internal capsule
GRE: gradient echo
MCP: middle cerebellar peduncle
MPRAGE: magnetization prepared rapid gradient echo
MRI: magnetic resonance imaging
NHP: non-human primate
NMT: NIMH macaque template
OT: optic tract
Put: putamen
QSM: quantitative susceptibility mapping
ROI: region of interest
SARM: subcortical atlas of the rhesus macaque
SN: substantia nigra
SPACE: sampling perfection with application optimized contrasts using different flip angle evolution
STN: subthalamic nucleus
T1w: T1-weighted
T2w: T2-weighted
TE: echo time
TR: repetition time
3D: three-dimensional
VP: ventral pallidum

## Notes

*Conflict of interest*: All authors claim that there are no conflicts of interest.

### Competing Interest Statement

The authors have declared no competing interest.

